# Incomplete lineage sorting of segmental duplications defines the human chromosome 2 fusion site early during African great ape speciation

**DOI:** 10.1101/2024.12.12.628057

**Authors:** Xinrui Jiang, Lu Zhang, Zikun Yang, Xiangyu Yang, Kaiyue Ma, DongAhn Yoo, Yong Lu, Shilong Zhang, Jieyi Chen, Yanhong Nie, Xinyan Bian, Junmin Han, Lianting Fu, Juan Zhang, Guojie Zhang, Qiang Sun, Evan E. Eichler, Yafei Mao

## Abstract

All great apes differ karyotypically from humans due to the fusion of chromosomes 2a and 2b, resulting in human chromosome 2. Yet, the structure, function, and evolutionary history of the genomic regions associated with this fusion remain poorly understood. Here, we analyze finished telomere-to-telomere chromosomes in great apes and macaques to show that the fusion was associated with multiple pericentric inversions, segmental duplications (SDs), and the rapid turnover of subterminal repetitive DNA. We characterized the fusion site at single-base-pair resolution and identified three distinct SDs that originated more than 5 million years ago. These three distinct SDs were differentially distributed among African great apes as a result of incomplete lineage sorting (ILS) and lineage-specific duplication. Most conspicuously, one of these SDs shares homology to a hypomethylated SD spacer sequence present in hundreds of copies in the subterminal heterochromatin of chimpanzees and bonobos. The fusion in human was accompanied by a systematic degradation of the three divergent α-satellite arrays representing the ancestral centromere creating five distinct structural haplotypes in humans. CRISPR/Cas9-mediated depletion of the fusion site in human cell lines significantly alters the expression of 108 genes, indicating a potential regulatory consequence to this human-specific karyotypic change.

## INTRODUCTION

Karyotype evolution is a critical aspect of evolutionary biology because it has been associated with speciation, adaptation, and disease^1–7^. In 1935, Wilson and Painter first observed that chromosome fusion could lead to speciation in flies^7^. As the field advanced, numerous instances of chromosome fusion have been documented in plant and animal speciation^1–4,8–10^. Several molecular mechanisms have been proposed as key drivers of chromosome fusion in speciation and oncogenesis, including telomere-telomere, telomere-centromere, tandem repeat, segmental duplication (SD), and retrotransposon expansions and fusions^11–15^.

Recent advances in genome editing and synthetic biology have provided deeper insights into the role of chromosome fusion in evolution^16–18^. Boeke and colleagues, for example, demonstrated that chromosome fusion could induce reproductive isolation in yeast, highlighting the potential of these genomic changes to influence speciation^19,20^. Additionally, studies on chromosome engineering in mice reveal significant chromatin conformation alterations in the chromosome fusion regions, further emphasizing the impact of karyotype evolution on genetic and phenotypic diversity^21,22^.

Human chromosome 2 (chr2) was formed from the fusion of nonhuman primate (NHP) chr2a and chr2b^23–25^, representing arguably the most significant karyotypic differences between humans and NHPs. Ijdo and Wienberg were the first to identify the fusion site at human chromosome 2q13-2q14 using cytogenetic banding approaches^25^. Subsequent studies uncovered dispersed SDs associated with this fusion site^26,27^. These analyses relied on partial genome assemblies generated using short-read technologies or Sanger sequencing of bacterial artificial chromosome (BAC) clones, often resulting in incomplete genetic information at the fusion site^26–29^. Consequently, the structure, evolutionary history, and function of the fusion site have remained only partially understood.

Available genomic assemblies and FISH experiments have shown an abundance of satellite sequences in the subtelomeric repetitive regions of NHP chr2a and chr2b, whereas such sequences are absent in humans^26,30,31^. During the fusion of human chr2, one of the centromeres in the fused chromosome becomes inactive, and degraded with independent transposable element (TE) retrotransposition events occurring at the degenerate site^32^. In addition, there has been considerable debate regarding the timing of the chr2 fusion event. Based on SD and SVA (SINE-VNTR-Alu) divergence, the fusion event was estimated to have occurred early in human evolution, 5-7 million years ago (mya) and 2.5-4.5 mya, respectively^26,33^. However, a recent study, utilizing clustered substitution statistics, proposed a much more recent origin of approximately 0.9 mya^34^.

The complex repetitive structure at the fusion site, subtelomeric repetitive regions, as well as the inactive centromere have made it challenging to reconstruct the evolutionary history and potential functional consequences of the fusion event. Here, we leverage the complete sequences of great apes^35,36^ and a macaque^37^ to revisit the structure and evolutionary history of human chr2. We aim to: (1) characterize the fusion site at a single-base-pair resolution, (2) study this in the context of epigenetic and structural changes occurring at the subtelomeric repetitive regions and degenerate centromere site, (3) use these data to create a model for human chr2 evolution, and (4) examine the potential functional consequences by creating fusion-site depletion cell lines.

## RESULTS

### Comparative sequence analysis of the human chromosome 2 fusion site

We performed a comparative analysis of 13 finished nonhuman great ape chromosomes^35^ and the finished macaque genome^37^ to human chr2 (Figure 1a, Methods). We identified numerous non-syntenic segments (76-154 regions ≥10 kbp in length) between humans and NHPs, accounting for 9.86 Mbp (macaque) and up to 57.5 Mbp (gorilla) of unalignable chromosomal sequence per species. Most of this sequence corresponded to various classes of repetitive DNA (Figure 1a, Supplementary Table 1), including satellite repeats, tandem and interspersed SDs. In particular, among the nonhuman African great apes, both chr2a and chr2b are acrocentric with the presence of subterminal heterochromatic caps (9.6-19.6 Mbp) demarcating the ends of the chromosomes and are composed of a 32 bp AT-rich satellite DNA (pCht) and SD spacer regions^26,31,35,38^. In addition, there are multiple pericentric and paracentric evolutionary inversions distinguishing the ape lineages from each other^39^, several of which share homology to the SD flanking regions of the fusion site (Figure 1b and Figure 2a). One pericentric inversion occurred in the common ancestor of African great apes after divergence from orangutans (chr2b) while another occurred before the *Pan*–gorilla split (chr2a)^39,40^ (Figure 1a). Furthermore, we identified a truncated SD containing the partial *CBWD2* at the chr2a pericentric inversion breakpoints, and another truncated SD with *FOXD4L1* at the chr2b breakpoint (Supplementary Figure 1)^28,29^.

**Figure 1.**
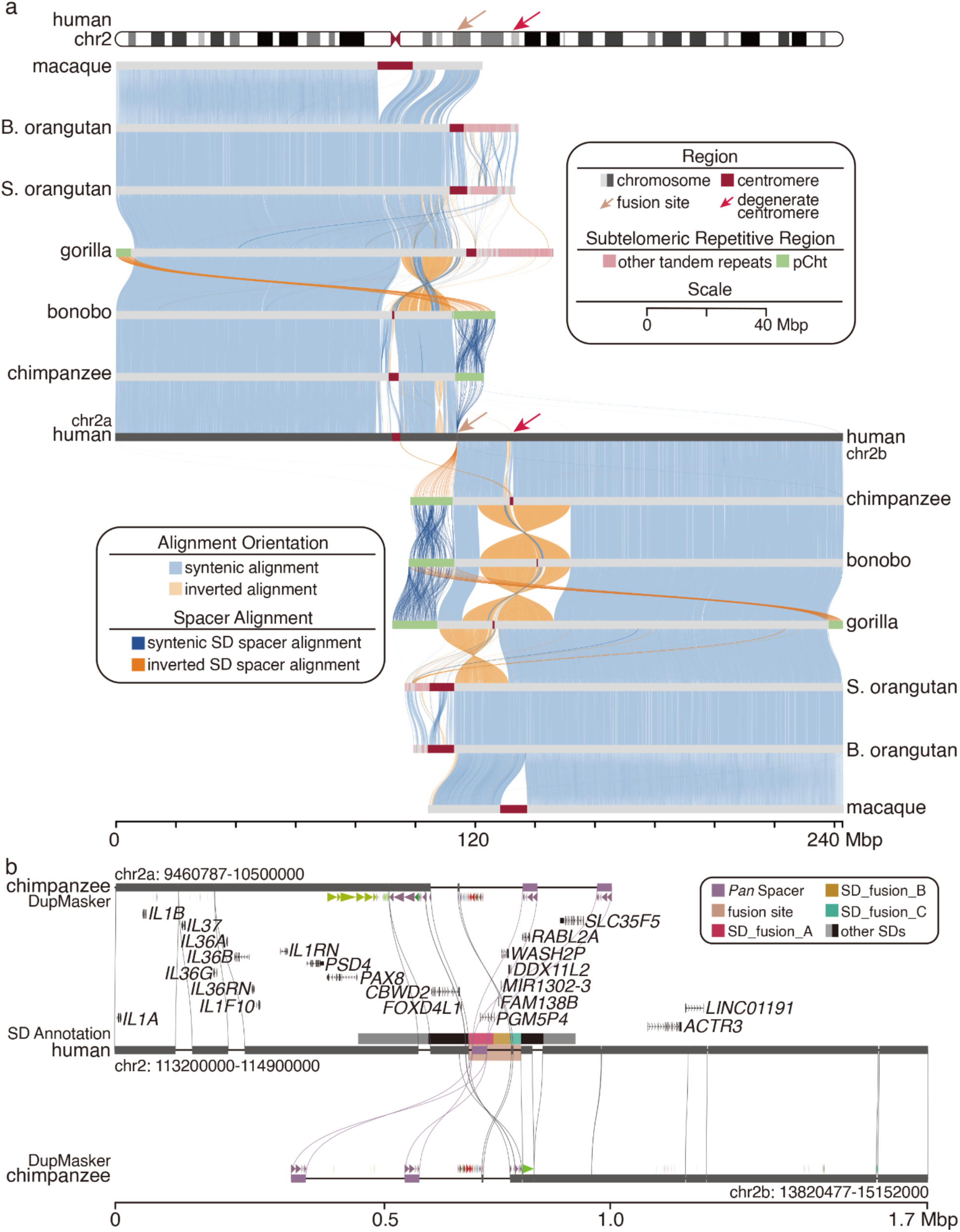
The comparative sequencing analysis of primate chromosome 2. (a) The syntenic comparison of chromosome 2 highlights the extent of primate chromosome 2 evolutionary rearrangements. Syntenic regions conserved in order (blue) are contrasted with evolutionary inversions (orange) and non-syntenic regions (breaks) corresponding to alpha satellite (dark brown), pCht subterminal satellite (green), and other satellite DNA (pink). B. orangutan and S. orangutan represent Bornean orangutan and Sumatran orangutan, respectively. (b) High- resolution analysis of the human fusion site (chr2:113,940,058-114,049,496, colored in amber) shows the non-syntenic breakpoint region in the context of annotated human protein-coding genes and segmental duplications (SDs). *Pan* SD spacers (purple blocks) in chimpanzee pCht show the homology of the partial region of human fusion site.

**Figure 2.**
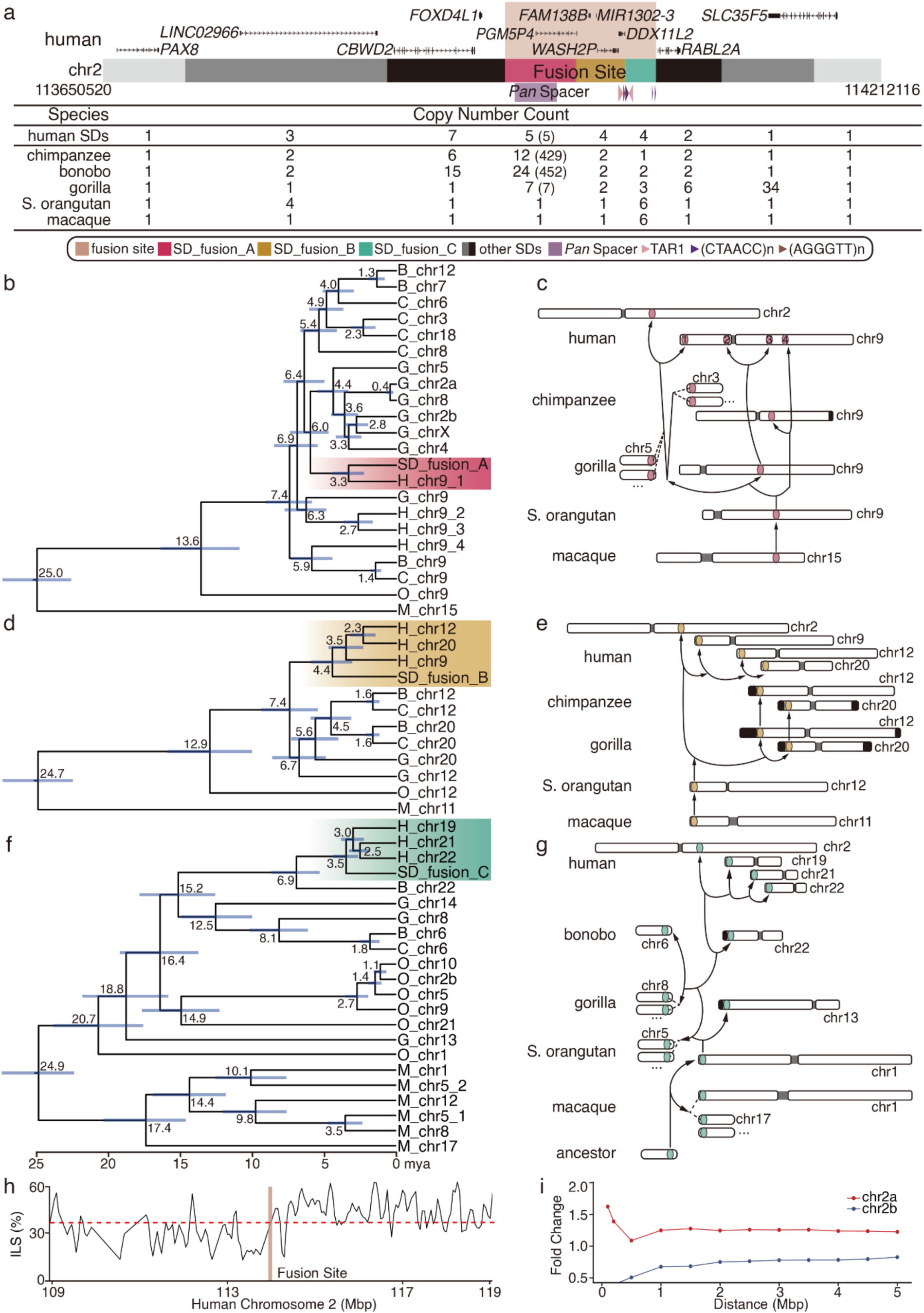
**The genomic structure and evolutionary history of SDs at the human fusion site.(a) Human chromosome 2 fusion site and SD organization**. A human genomic segment (chr2:113,650,520-114,212,116) with gene annotations shows the region where chromosome 2a and 2b fused (chr2:113,940,058-114,049,496, amber). The region consists of a large (∼455 kbp) duplication block made up of seven SDs with variable copy numbers in each ape genome. The three SDs at the fusion site are designated as SD_fusion_A, SD_fusion_B, and SD_fusion_C. The table indicates the copy number of the full-length homologous segments for each primate genome (with brackets indicating the SD_fusion_A copy number of the derived sequence in the subterminal heterochromatic caps of chimpanzees and bonobo; see Supplementary Table 2 for the detailed alignments at the fusion site). (b-g) Phylogenetic trees based on a (b) 53 kbp (SD_fusion_A), (d) 68 kbp (SD_fusion_B), and (f) 23 kbp (SD_fusion_C) multiple sequence alignment show that these regions have been subject to incomplete lineage sorting (ILS). The chromosome schematic depicts the genomic locations of SD_fusion_A (c), SD_fusion_B (e), and SD_fusion_C (g) in each primate. (h) Proportion of human–gorilla and *Pan*–gorilla tree topology in a 500 bp window of 5 Mbp flanking region near the fusion site. The red dotted line represents the mean value of the whole chr2. (i) Fold change of the mean proportion of discordant topologies in the flanking region compared with the whole genome average. (Note, chromosome names in the great apes refer to the human homologous chromosome also known as the phylogenetic group designation.)

### Evolutionary reconstruction of the ancestral fusion site

We further compared human chr2 with *Pan* chr2a and chr2b to precisely characterize the ∼109 kbp fusion site in the human T2T- CHM13v2.0 genome assembly (2q14.1, chr2:113,940,058-114,049,496) (Figure 1b and Supplementary Figure 2). Leveraging data from 47 human genomes from the Human Pangenome Reference Consortium (HPRC)^41^, we confirmed that there is a complete absence of large structural variants within this fusion site in 94 sequence-resolved human haplotypes, suggesting a fixed structural haplotype for the entire fusion site (Supplementary Figure 3). Neither nucleotide diversity (π) nor Tajima’s D values show significant reductions at the fusion site in African or non-African populations, relative to the chr2 average (Supplementary Figure 4) consistent with the locus evolving neutrally.

Within the larger ∼455 kbp duplication block (seven independent SD blocks) demarcating the fusion site region, we narrowed the fusion site to a ∼109 kbp segment consisting of three distinct, high-identity SDs (≥98% identity and ≥20 kbp in length) (Figure 1b, Figure 2a, Supplementary Figures 5-12, and Supplementary Tables 2-3). In humans, these SDs share homology with sequences corresponding to chr9 (SD_fusion_A, 50 kbp), chr12/20 (SD_fusion_B, 36 kbp), and chr22 (SD_fusion_C, 22 kbp) (Figure 2a, Supplementary Table 2). Among African apes, each SD is highly variable in copy number showing evidence of shared ancestral locations as well as lineage-specific duplications (Supplementary Table 2). Using macaque and orangutan as outgroups, we reconstructed the evolutionary history of each SD estimating the divergence timepoint from each SD to its nearest nonhuman great ape neighbor (Figure 2b-g, Methods).

SD_fusion_A (50 kbp) is present as a single copy in macaques and orangutans and corresponds to a partial duplication of *PGM5*^29^ (14 exons of an ancestral gene), a gene involved in carbohydrate metabolism, originating from African great ape ancestral chr9 (phylogenetic group IX) (Figure 2b-c). This SD began to duplicate in an interspersed configuration in the common ancestral lineage of African great apes (∼7.4 mya) with the locus expanding to highest copy number in chimpanzees and bonobos where it defines the SD spacer region demarcating large blocks of pCht on chr2a and chr2b and other chromosomes (chr3, chr5-chr13, chr15-chr22, and chrX in chimpanzee; each chromosome in the bonobo genome) (Figure 2b-c and Supplementary Figure 6). Of note, the SD_fusion_A (chr2) and other human copies share a monophyletic origin (∼3.3 mya) most closely related to gorilla copies ∼6 mya (95% CI: 4.8-7.4 mya) instead of chimpanzee or bonobo—a pattern consistent with incomplete lineage sorting (ILS).

Similarly, SD_fusion_B, is a 36 kbp segment corresponding to the partially truncated *WASH2*^42^ (11 exons of an ancestral gene that potentially regulates actin cytoskeleton dynamics), *FAM138B* (3 exons of an ancestral gene of unknown function), and *DDX11* (3 exons of an ancestral gene implicated in DNA metabolism). The segment exists as a single copy in both macaques and orangutans and originated from a locus mapping to ancestral African great ape chr12 (phylogenetic group XII) (Figure 2d-e and Supplementary Figure 7). All human copies mapping to human chr2, chr20, chr12, and chr9 show a monophyletic origin (∼4.4 mya, 95% CI: 3.1-5.8 mya) suggesting human-specific duplications or interlocus gene conversion events^43^. With respect to nonhuman African great apes, however, this clade diverges from a distinct monophyletic clade that includes chimpanzee, gorilla, and bonobo (∼7.4 mya, 95% CI: 5.4-9.3 mya) (Figure 2d). This topology is once again consistent with ILS.

In contrast to SD_fusion_A and SD_fusion_B, SD_fusion_C (the most distal, 22 kbp) shows evidence of independent duplication in all primates, including macaques and orangutans, with the exception of chimpanzees where it exists as a single copy on chimpanzee chr6 (Supplementary Figure 8). All human SDs show a monophyletic origin (∼3.5 mya, 95% CI: 2.7- 4.3 mya) but show a deep coalescence with another genomic segment present only on bonobo chr22 (phylogenetic group XXII), dating back to ∼6.9 mya (95% CI: 5.3-8.6 mya) (Figure 2f). Notably, SD_fusion_C and its flanking region correspond to a larger SD that aligns exclusively between the human chr2 SD_fusion_C site and bonobo chr22, but not with any other NHPs (Supplementary Figure 13). Thus, SD_fusion_C and its flanking region likely represent an ancestral genomic structure in the latest common ancestor (LCA) of African great apes, which subsequently sorted in the human and bonobo lineages.

We extended the ILS analysis to the 5 Mbp mapping proximally (chr2a) and distally (chr2b) to the fusion site in humans (Methods) using a 500 bp windowed approach^44^ (Supplementary Table 4). We observe a sharp transition in the proportion of ILS windows at the site of the chr2 fusion. Specifically, we find the proportion of ILS rises to 45.6% (a 1.39-fold excess compared to the chr2 average) as we approach the fusion site distally (chr2b side). In contrast, the proximal portion appears depleted for ILS segments (Figure 2h-i, Supplementary Figure 14 and Supplementary Table 5). There is, thus, a polarized pattern of ILS with maxima and minima occurring on either side of the fusion site.

### The refined analysis of telomeric sequences at the fusion site

Previous investigations identified inverted telomeric sequences (TTAGGG/CCCTAA) at the fusion site, suggesting a telomere-to-telomere (T2T) fusion^25^. We confirmed the presence of interstitial telomeric sequences (CCCTAA, chr2: 114,027,659-114,028,207) at SD_fusion_C and its corresponding SD on human chr22 (TTAGGG, chr22:51,254,086-51,323,279) and similar sequences in bonobo chr22 (TGAGGG, chr22: 60,722,979-60,724,048). The sequence (GGGTTA) was identified at SD_fusion_B (chr2: 114,027,333-114,027,658) and its corresponding SD on human chr12 (TAACCC, chr12:2,843-3,030) and bonobo chr20 (TAACCC, chr20:1,139,021-1,139,266), but not at its corresponding SD on human chr20 (Supplementary Figure 15). These observations argue that these telomeric sequences were present on SD_fusion_B and SD_fusion_C prior to the fusion event. In addition, two telomeric-associated repeats (TAR1) flank the telomeric sequences of SD_fusion_B and SD_fusion_C^27^. Given the genomic structure of TAR1 and telomeric sequences on these SDs (Supplementary Figures 15-16), we propose that, in the ancestral configuration, each SD contained a TAR1 element and telomeric sequences, arranged in an inverted orientation on ancestral chr2a and chr2b. The end-to-end fusion of these two distinct SDs may have facilitated the fusion of ancestral chr2a and chr2b, leading to the vestigial presence of two TAR1 elements and the (CTAACC) and (GGGTTA) repeat motifs at the human fusion site.

### African great ape subterminal satellite repeat expansion and human chromosome 2 fusion

Reconstructing the evolutionary history of the fusion event has been challenging due to extensive SD and lineage-specific turnover of the subterminal heterochromatic caps in *Pan* and gorilla^35^ (Supplementary Table 6). With the exception of the short arm of gorilla chr2a, the corresponding regions in both *Pan* and gorilla are composed of nearly continuous megabase-pair tracts of satellite DNA interspersed with SD spacers^35^. The satellite sequence is made of a tandem 32 bp repeat motif (pCht)^31^ punctuated on average every 287 kbp in *Pan* and every 389 kbp in gorilla by a SD spacer (Figures 3a, Supplementary Figures 17-19). The interrupting SD spacers are variable with a modal length of 32 kbp in the *Pan* lineage and 33.7 kbp in gorilla (Supplementary Figure 17) and correspond to hypomethylated pockets flanked by the hypermethylated satellite DNA^35^ (Figure 3a). Although both the gorilla and chimpanzee SD spacers differ in sequence composition and the subterminal heterochromatic cap is largely thought to have evolved independently in both lineages^26,35^, the net effect is that the subterminal portions of both chimpanzee chr2a and chr2b share 23.6 Mbp of high-identity sequence homology involving both the SD spacers and pCht satellite DNA. In the case of the gorilla, the homology is restricted to the pCht satellite DNA for both arms of gorilla chr2b and the q-arm of gorilla chr2a. The organization of the p-arm of gorilla chr2a differs considerably and is much more similar to the organization found in orangutan, which is enriched in HSatIII-like repeat sequences (Figure 3b). It is classified as an acrocentric short arm lacking a nuclear organizing region^35^.

**Figure 3.**
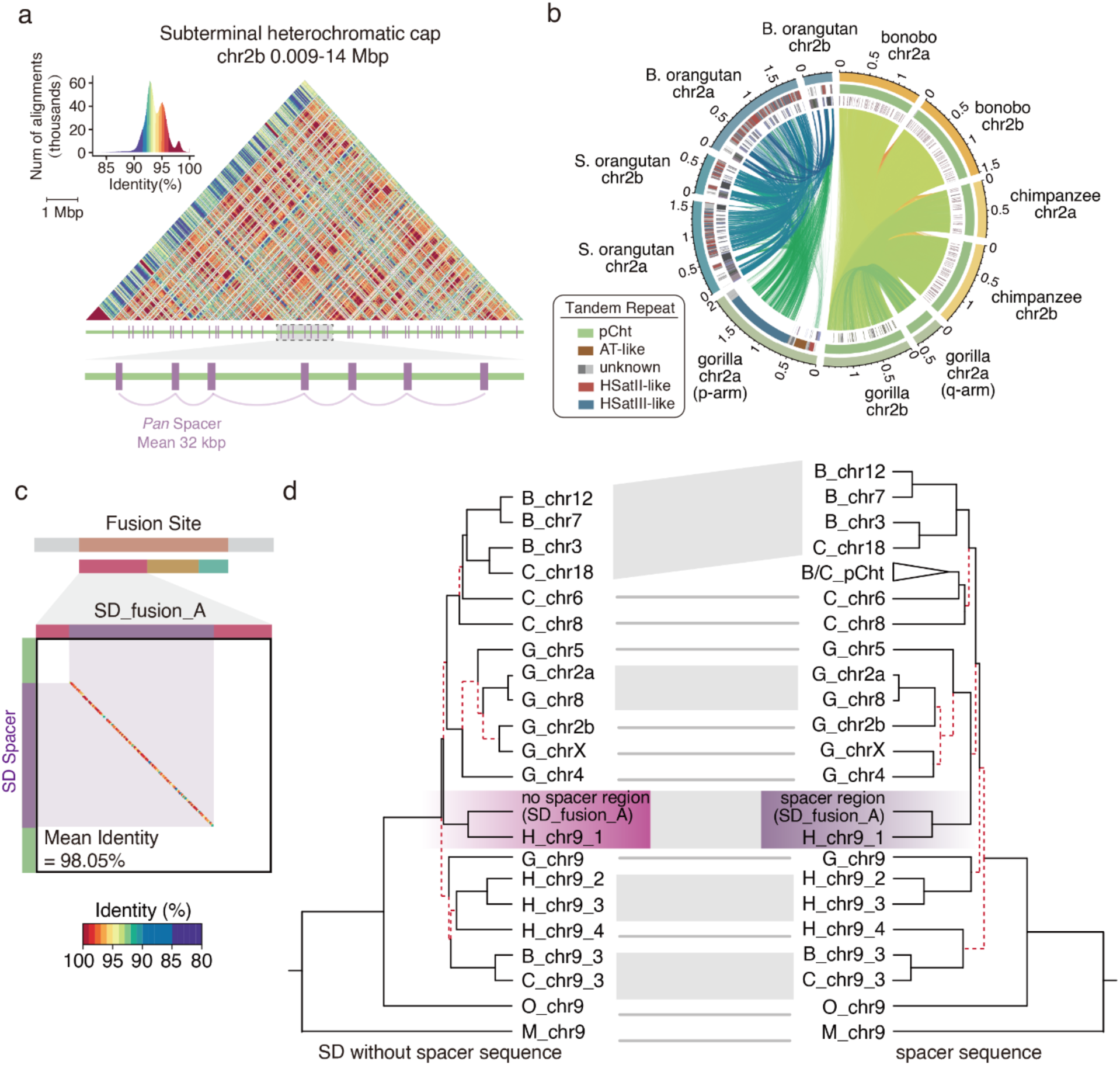
Rapid turnover of subtelomeric repetitive regions in primates and evolutionary connection between SD spacers in *Pan* lineage and SD_fusion_A at the fusion site. (a) The identity heatmap of the subterminal heterochromatic cap for the p-arm of chimpanzee chr2b. SD spacer elements (purple) are annotated below and are flanked by large tracts of pCht satellite (green) (zoomed-in panel shows the structure at higher resolution). (b) Circos plot shows sequence identity of chr2a and chr2b among the apes highlighting the rapid turnover of satellite DNA. The three layers (outer to inner) represent the subtelomeric region, tandem repeat satellites, and transposon element annotations. (c) Dot plot shows the synteny between a bonobo SD spacer within the heterochromatic cap vs. human SD_fusion_A segment. The sequence identity was calculated at a 100 bp resolution, with the dot color representing sequence identity ranging from 80% (purple) to 100% (red). (d) Phylogenetic topology comparison shows nearly consistent ILS evolutionary pattern of SD_fusion_A (excluding spacer sequence) and SD spacer. Each node is supported by a bootstrap value of 100/100. The red lines indicate discordant tree topologies within each major clade between the two trees.

Importantly, the SD spacer that expanded in the subterminal heterochromatic caps in chimpanzee and bonobo shares 98.5% identity with the SD_fusion_A segment mapping at the fusion site on human chr2 (Figure 3c). All 429∼452 SD spacers within the *Pan* pCht regions, including those at the ends of these chromosomal regions, are monophyletic in origin estimated to have expanded approximately ∼5.5 mya (95% CI: 4.2-6.8 mya). This subterminal SD spacer (32 kbp) in *Pan* is 18 kbp smaller than SD_fusion_A (50 kbp) where the shared LCA predates the human–*Pan*– gorilla divergence (Figure 2b and 2c). Comparing the phylogenies of the sequence unique to SD_fusion_A and sequence shared with the subterminal heterochromatic cap SD spacers shows nearly coincident ILS topology (generalized Robinson–Foulds distance=0.28, p=1.6×10^-4^). This indicates that the SD spacers within *Pan* heterochromatic caps are derived from the duplicated sequence that gave rise to ancestral SD_fusion_A. Thus, the hyperexpansion of subterminal satellite DNA associated with subterminal heterochromatic caps in *Pan* and the chr2 fusion are linked genetically with two different evolutionary trajectories and karyotypic consequences in human and *Pan* (Figure 2c-d, Supplementary Figure 20 and Supplementary Table 7).

### Centromere retention and degeneration in human chromosome 2

An important consequence of the human chr2 fusion was that the ancestral chr2b centromere in NHPs became inactive in our human lineage (Figure 4a). Chr2a differs from chr2b by the presence of large tracts of HSatII arrays in humans and HSatIII arrays in *Pan* (Supplementary Figures 21-22). Overall, the active human centromere in humans is more similar to that of gorilla with respect to suprachromosomal family (SF) organization. Both humans and gorillas possess SF2, whereas chimpanzees and bonobos have SF3.

**Figure 4.**
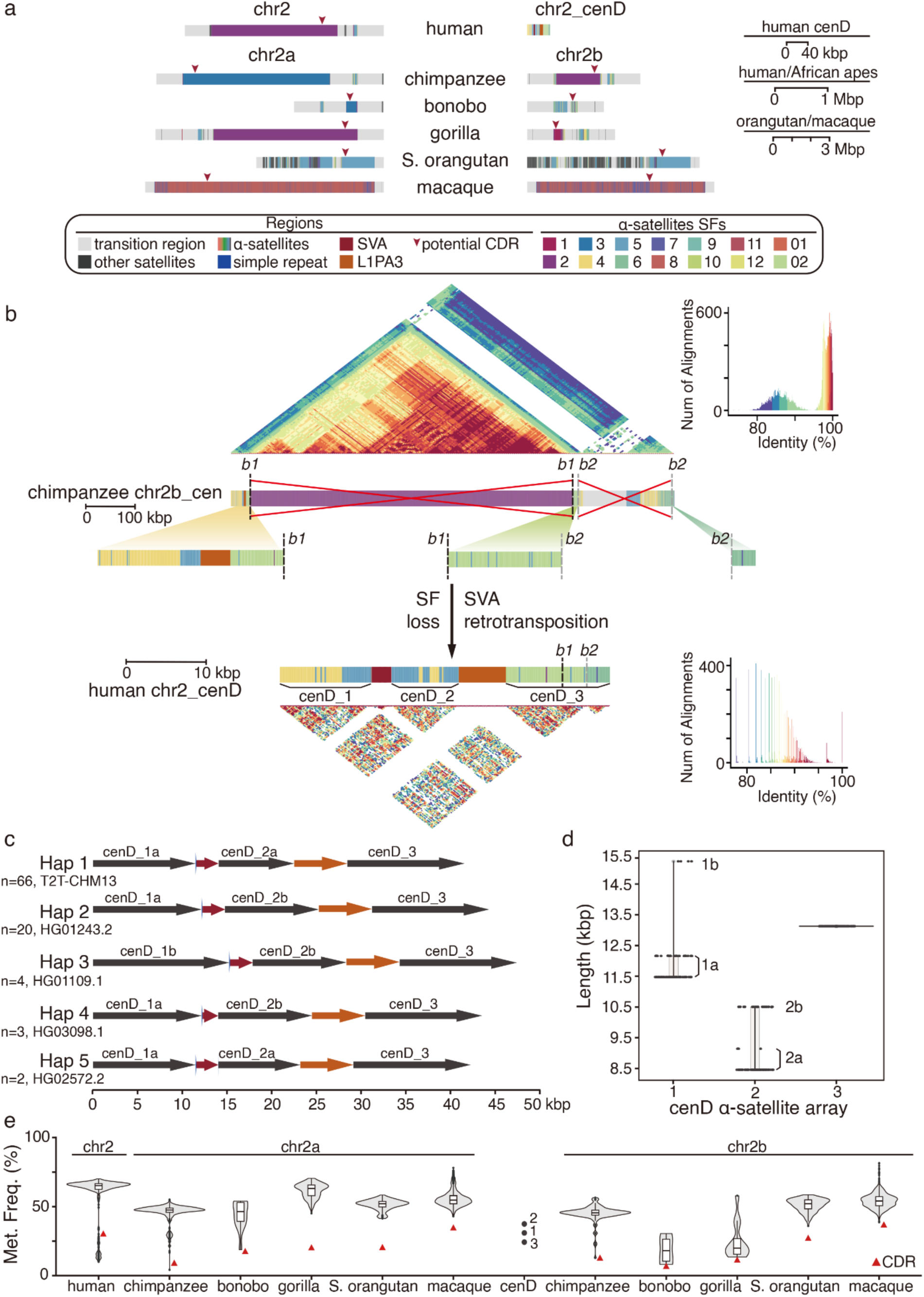
**Comparative analysis of active and inactive centromeric regions of human chromosome 2**. (a) Genomic structure and suprachromosomal family (SF) annotation of centromeres in human chr2, degenerate site, and NHP chr2a/chr2b. Different SFs are shown by various colors, with centromere dip regions (CDRs) marked by arrows. (b) Comparison of the centromere in chimpanzee chr2b with the human degenerate centromeric region. The middle panel shows the SF annotations for the chimpanzee centromere and human centromere degenerate region. Potential α-satellite deletion breakpoints (*b1* and *b2*) are indicated with dotted lines. The heatmaps of each region are shown in upper and lower, respectively. (c) The panel illustrates five distinct structural haplotypes of the degenerate centromere site in humans. (d) The panel shows the lengths of three α-satellite arrays in humans. (e) The methylation degree of HOR across NHPs and of three α-satellite arrays at the degenerate centromere site. The red triangle represents the methylation frequency of CDR in each chromosome (Met. Freq. strands for methylation frequency).

We compared the active chromosome centromere of chimpanzee chr2b with the structure of the vestigial centromere in humans. We identified three distinct α-satellite arrays (chr2:132,644,386– 132,685,996) in humans^45^ with homology to chimpanzee that had been interrupted by various simple repeats (CCTCTC) and retrotransposon elements (SVA and L1PA3) in the human lineage (Figure 4b). All three satellite arrays were derived from divergent monomeric α-satellite regions ancestral to human and chimpanzee, rather than from higher-order repeats (HORs). Our analysis of 94 human genome assemblies^39^ identifies five distinct structural haplotypes due primarily to length variation of the first two α-satellite arrays (Figure 4c-d and Supplementary Table 8). A sequence comparison using unique *k*-mers from this region shows that ∼97.5% (8,246/8,460) and ∼96.7% (8,181/8,460) are also identified in Neanderthal and Denisovan genomes, respectively (Methods), confirming that the centromeric degeneration occurred long before the divergence of modern and archaic humans^32^.

We also compared methylation patterns of the NHP chr2a and chr2b centromeres (Figure 4e). The human α-satellite arrays mapping to the degenerate site show significantly lower methylation levels, when compared to typical NHP HORs (excluding centromere dip regions [CDRs], p=0.02), with the exception of bonobo chr2b and gorilla chr2b. HORs in bonobo chr2b and gorilla chr2b are significantly hypomethylated when compared to other NHP HORs (p=1.92×10^-6^), possibly due to the smaller size of these HORs (lengths: 71 kbp for bonobo chr2b, and 106 kbp for gorilla chr2b).

### Functional assessment of the fusion site by depletion

Four putative noncoding genes/pseudogenes (*PGM5P4*, *FAM138B*, *WASH2P*, and *DDX11L2*) have been annotated at the site of the human chr2 fusion. According to GTEx short-read RNA sequencing (RNA-seq), these four genes/pseudogenes are expressed in testis, esophagus, fallopian tube, and cerebellum tissues^46^ (Supplementary Figure 23). Further, both *PGM5P4* and *WASH2P* are supported by long-read Iso-Seq transcript data from CHM13hTERT^47^ and kidney tissue from ENCODE^48^. In addition, methylation analysis demarcates a prominent CpG island showing the promoters/enhancers of *PGM5P4*, as identified using ONT reads from the T2T-CHM13 cell line^47^ (Supplementary Figure 24).

To explore the potential function of the fusion site, we used CRISPR/Cas9 to delete this region in a kidney-derived epithelial-like cell line (HEK293T) (Figure 5a). Two independent pairs of single guide RNA (sgRNA; L-sg1 and R-sg; L-sg2 and R-sg) were designed for the depletion (Figure 5b), and the heterozygous depletion of the fusion site in both sgRNA groups was confirmed by PCR and Sanger sequencing (Figure 5c and Supplementary Table 9). We then conducted RNA-seq, generating 53.28 Gbp, 67.42 Gbp, and 122.41 Gbp of data for the L-sg1 depletion, L-sg2 depletion, and wild-type (WT) cell lines (three biological repeats for each depletion cell line and six biological repeats for WT cell lines) (Supplementary Table 10).

**Figure 5.**
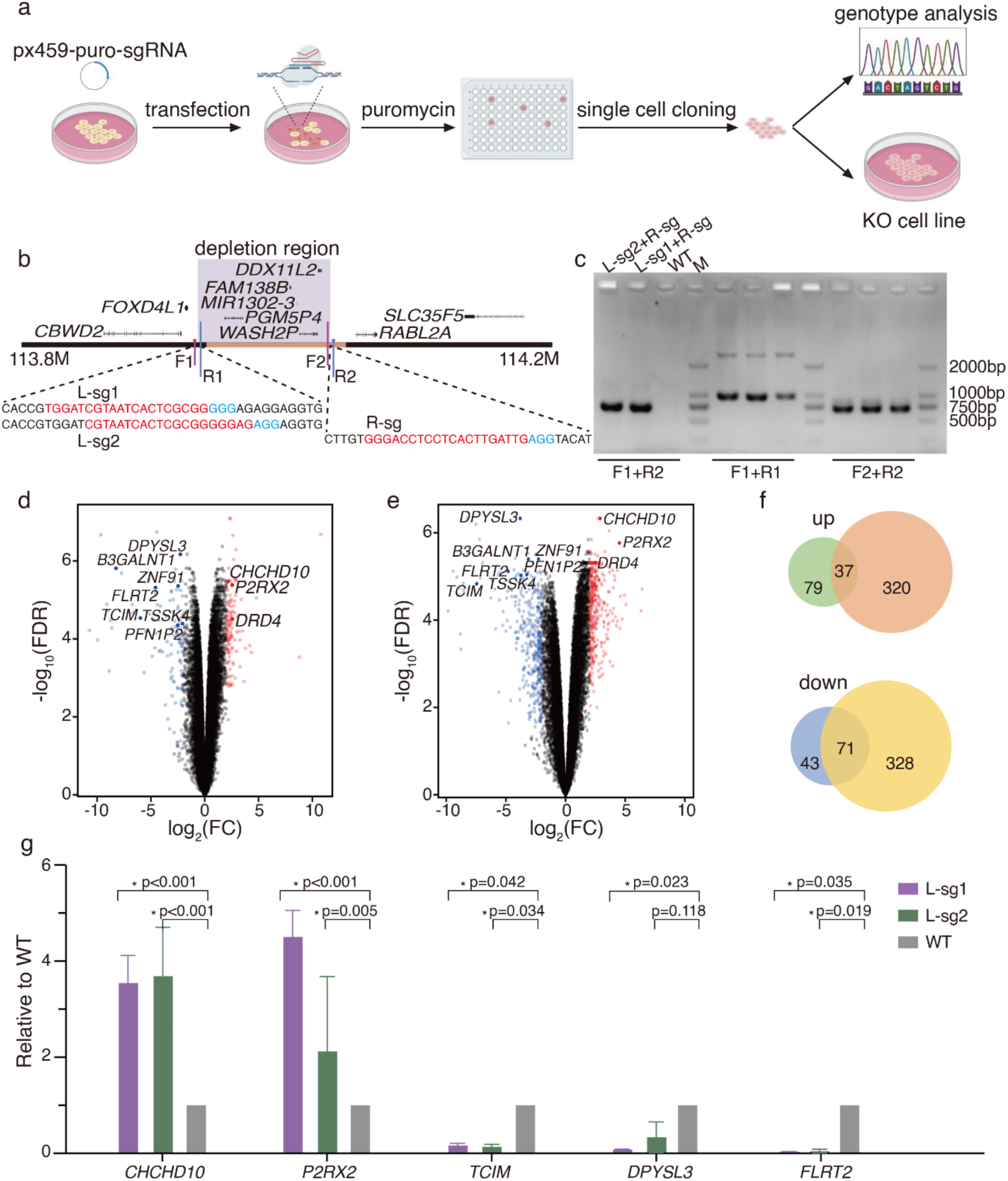
**Fusion site knockout and gene expression alteration**. (a) Schematic representation of the depletion experiments performed in HEK293T cell lines. (b) Two distinct sgRNA pairs (L-sg1 and R-sg; L-sg2 and R-sg) were designed for the depletion experiments (left panel). Red sequences indicate sgRNA, while blue sequences indicate PAM sites. (c) The PCR analysis confirms heterozygous depletion in cell lines. WT represents the wild-type cell lines. (d) and (e) Differentially expressed genes (DEGs) are shown in volcano plots (left panel for the L-sg1 and R-sg pair, right panel for the L-sg2 and R-sg pair). Genes with log2 fold change (log2FC) values >2 and FDR <0.05 are highlighted in red (upregulated) and blue (downregulated). The overlap of top 10 DEGs in each group are emphasized in deeper red/blue, with gene names indicated. (f) Venn diagrams illustrate the overlap of upregulated and downregulated genes between both depletion groups. (g) The RT-PCR analyses confirmed 5 of 10 top-rank DEGs. WT represents the relative gene expression levels normalized to *GAPDH* in wild-type cell lines.

Differential gene expression analysis identified 37 upregulated and 71 downregulated genes in both depletion cell lines (Supplementary Figure 25 and Supplementary Table 11, Methods).

These genes were significantly enriched in pathways related to the regulation of transcription by RNA polymerase II (p=2.2×10^-3^). Among the top ten differentially expressed genes (DEGs) in both depletion cell lines, three were upregulated and seven were downregulated (Figure 5d-e).

We further validated five of ten gene expression changes using RT-PCR in the depletion cell lines (Figure 5g and Supplementary Table 12).

## DISCUSSION

Complete T2T sequence for chromosome 2, 2a, and 2b among the great apes^35–37^ allowed us to systematically examine the complex genomic architecture and further refine the evolutionary history of the human-specific chr2 fusion event. There are three important conclusions. First, the fusion event was intimately associated with SDs that have been restructuring genomes and chromosomes throughout great ape evolution^26,28^. The 109 kbp fusion site in humans consists of three independent SDs, each with distinct trajectories in different ape lineages, and these are further embedded in a larger duplication block of ∼455 kbp. All SDs are highly variable in copy number among the great apes, and many have been reused as breakpoint sequences during great ape evolution. Although cytogenetic studies previously reported two pericentric inversions on ancestral chr2a and chr2b^26^, we observe that the flanking region at the human fusion site (chr2a, proximal side) shares 98.4% identity with an SD flanking the pericentric inversion that distinguishes Sumatran orangutan chr2b from gorilla chr2b. Moreover, SD_fusion_A shares 97.8% identity with the SD spacer sequence of the *Pan* subterminal heterochromatic caps where it expanded to hundreds of copies in the chimpanzee and bonobo lineages but not in gorilla.

Second, we show that the fusion site has been strongly subjected to ILS. All three SDs, for example, show phylogenetic signatures consistent with ILS with a maximum occurring at the fusion site suggesting that the fusion event occurred >5 mya. Thus, the fusion event is not a recent evolutionary event but potentially occurred during African great ape speciation when the effective population size was predicted to be much larger than contemporary ape populations^35,44^. Given fossil evidence that *Australopithecus* existed around 2-4 mya, *Paranthropus* around 1-3 mya, and the earliest *Homo* fossils around 2-3 mya^49–51^, we speculate that this fusion event did arise in the *genus Homo* but rather occurred in ancestral great ape populations that would give rise to humans potentially by creating a stasipatric speciation barrier^52^.

Third, the fusion site was associated with extensive subtelomeric satellite turnover and epigenetic differences among great apes. The two satellite motifs present in the subtelomeric regions of gorilla chr2a and chr2b are distinct from each other: one resembles the subtelomeric repetitive regions of orangutans, while the other resembles the genomic architecture of *Pan* species. SD_fusion_A, in part, defines the chr2 fusion site but also associates with large tracts of hypermethylated pCht satellite repeats that define subterminal heterochromatin in chimpanzees and bonobos^35^ (Supplementary Figure 26). The juxtaposition of novel SDs at the fusion site and the shift from heterochromatic or acrocentric DNA at the termini of chr2a and chr2b led to methylation differences among the great apes at this locus. Indeed, our depletion experiments in humans show consistent gene expression changes suggesting that the euchromatization of the fusion site may have had more global genome-wide regulatory effects.

We propose two evolutionary scenarios for the formation of human chr2. In the first, the large effective population size of the human–*Pan*–gorilla ancestral population facilitated the coexistence of several NHP chr2a and chr2b subtelomeric structural haplotypes 5-7 mya. In one of these structural configurations, SD_fusion_A and SD_fusion_B became juxtaposed and duplicated to the subtelomeric region of human–*Pan*–gorilla ancestral chr2a prior to the fusion event (Figure 6a). Similarly, SD_fusion_C was duplicated to the subtelomeric region of ancestral chr2b (where the full-length structure is still retained subtelomerically in bonobo); yet, it is no longer located on other nonhuman great ape chr2b due to the rapid exchange of subtelomeric SDs and evolutionary turnover of satellite DNA^26,53^. Subsequently, the telomeric and TAR1 sequences in SD_fusion_B and SD_fusion_C mediated the fusion (Figure 6a). In the *Pan* and gorilla lineages, chr2a and chr2b (only in chimpanzee and bonobo) experienced a different evolutionary trajectory associated with the formation of subterminal heterochromatic caps.

**Figure 6.**
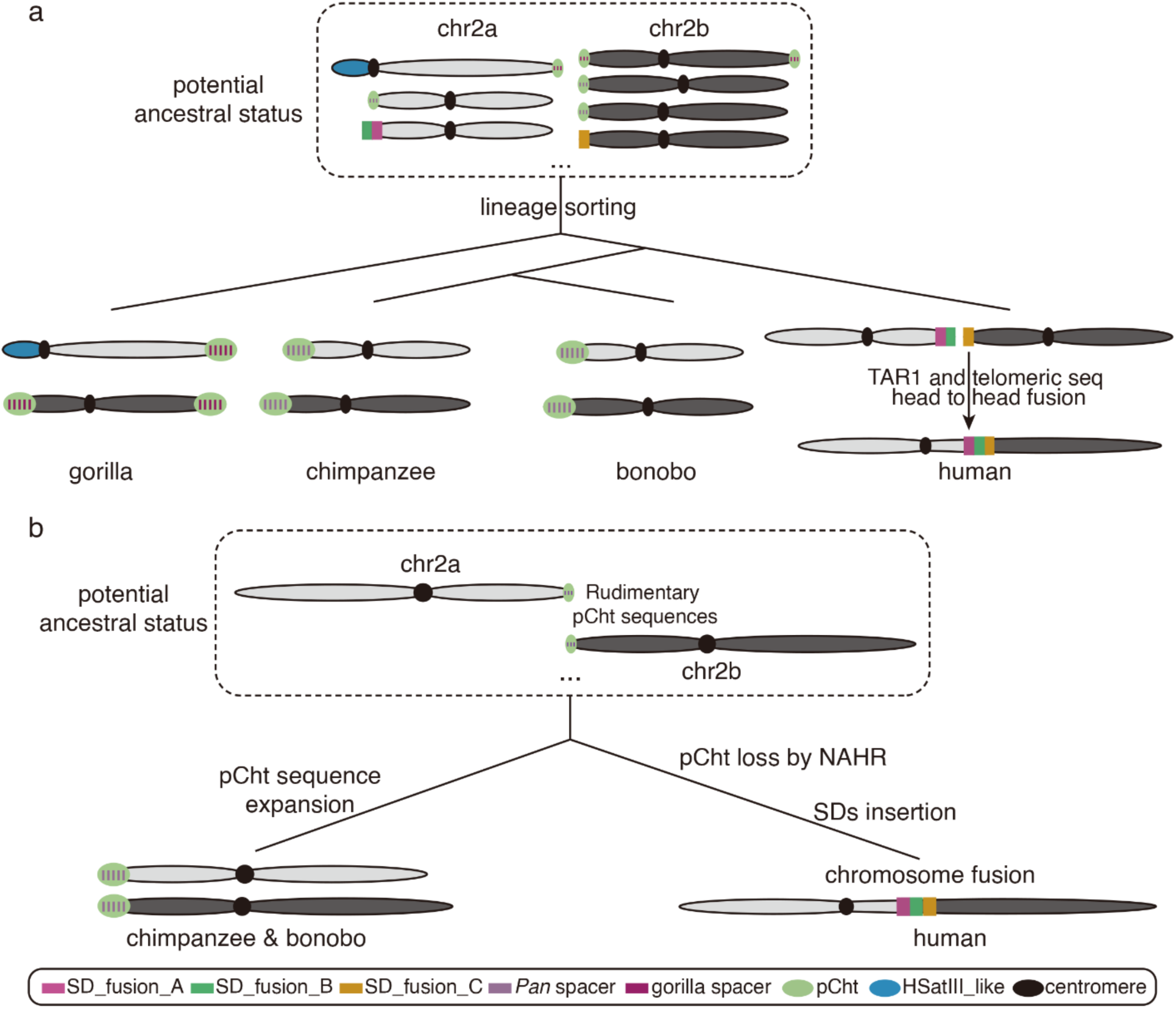
**Models for chromosome 2 evolution**. Two different models are depicted for the origin of the human chromosome 2 fusion. (a) In the last common ancestor of humans, bonobos, chimpanzees, and gorillas, diverse ancestral genomic structures emerged at the ends of chr2a and chr2b. In the human ancestral lineage, chr2a and chr2b consisted of complex SD blocks that emerged as a result of ILS, and then became juxtaposed by a telomere-to-telomere fusion. (b) Rudimentary pCht sequences were present in the ancestral lineage of humans and chimpanzees in association with SD_fusion_A. Nonallelic homologous recombination (NAHR) occurred between these in the human lineage eliminating pCht in humans but retaining SD_fusion_A with other SDs subsequently duplicating to the location. In chimpanzees, an independent expansion of the SD spacers and pCht sequences occurred leading to the formation of the subterminal heterochromatic caps.

Previous studies^26,35^ have shown that the heterochromatic caps likely evolved independently in both gorilla and chimpanzee, albeit convergently with a similar architectures where hundreds of kilobase pairs of satellite pCht DNA are punctuated by a ∼30 kbp SD spacer region that defines a pocket of hypomethylation. The SD spacers in gorilla and chimpanzee subterminal heterochromatin caps are distinct but in both chimpanzee and bonobo the SD spacers represent a derivative of the SD_fusion_A confirming that this sequence was present subtelomerically for chr2a in the human–*Pan* ancestor. SD_fusion_A subsequently expanded to hundreds of copies in chimpanzee and bonobo in conjunction with the subterminal heterochromatic satellite. Yet, in the human lineage, SD_fusion_A remains a low-copy SD with no trace of pCht satellite sequence in the human genome^35^.

Alternatively, the SDs evolved similarly, but in the common ancestor of human and *Pan,* there was an incipient association with rudimentary pCht satellite sequences with the SD_fusion_A sequence that mediated exchange between ancestral chr2a and chr2b via nonallelic homologous recombination (NAHR) or ectopic exchange. Subsequently, during NAHR between full-length SD_fusion_A elements, the pCht regions were lost in the human lineage and two additional SDs were inserted at the fusion breakpoint region (Figure 6b). In the *Pan* lineage, a portion of the SD_fusion_A became hyperexpanded defining the subterminal SD spacer region of all heterochromatic caps in bonobo and chimpanzee. In support of this model, it is known that in both chimpanzee and gorilla the subterminal satellite DNA forms unique post-bouquet structures in germ cells and are hotspots of ectopic exchange between nonhomologous chromosomes^54^. If such exchanges occurred at the edge of the incipient subterminal heterochromatic caps before there were many copies of the subterminal satellite DNA, it would help explain the absence of pCht sequence in the human genome—i.e., the fusion of chr2a and chr2b helped eliminate the potential for the formation of subterminal heterochromatic caps.

Irrespective of the model, there is evidence that SDs in the LCA and ILS potentially facilitated by a large effective population size played a key role^35,44^. These aspects helped maintain and diversify karyotypic structures of the ancestral chr2a and chr2b for possibly millions of years. The divergent fates of the subtelomeric repetitive regions of these ancestral chromosomes, e.g., pCht expansion and SD spacer insertions in the *Pan* lineage, or the SD insertions in the human fusion site, moreover, may have driven the speciation between humans and NHPs through ILS (Supplementary Figure 27). In this light, it is interesting that we document an asymmetric ILS pattern, with an increase in ILS segments on the distal side (chr2b) and a decrease on the proximal side (chr2a) of the fusion site. This suggests diverse evolutionary trajectories for the ancestral subtelomeric regions of NHP chr2a and chr2b highlighting the potential important role of both SDs and ILS in understanding chromosomal evolution, speciation, and sequence turnover.

## Methods

### Data resources and comparative analysis

The great ape and macaque T2T genome assemblies are publicly available (Data Availability). Syntenic relationships among primate chr2/chr2a/chr2b were assessed using minimap2^55^ (v2.24) with the following parameters: ‘-c -x asm20 --secondary=no --eqx -Y -K 8G -s 1000’ and visualized using SafFire (https://mrvollger.github.io/SafFire). We identified centromeres, transposons, and subtelomeric satellites based on RepeatMasker (v4.1.4) annotation (https://www.repeatmasker.org/).

### Fusion site characterization and population analysis

To precisely define the fusion site, we aligned NHP chr2a and chr2b to human chr2 using minimap2 (v2.24) with the following parameters ‘-x asm20 -r500,20000 -s 2000 -p 0.01 -N 1000 --cs’. The alignments were visualized using the minimiro (https://github.com/mrvollger/minimiro). We used VCFtools^56^ (v0.1.16) to calculate nucleotide diversity (pi) and Tajima’s D for population genetic analyses with the following parameters ‘--window-pi 20000 --window-pi-step 10000’ and ‘--TajimaD 20000’.

### SD annotations and phylogenetic analysis

SD annotations of the fusion site in the T2T- CHM13v2.0 were based on the UCSC SEDEF-SD track (https://genome.ucsc.edu/). In this study, we focused on the SDs with an identity of ≥98% and length ≥20 kbp. These SD sequences were aligned to NHP T2T genomes to identify homologous regions using minimap2 (v2.24) with the parameters: ‘-cx asm20 -p 0.5 --eqx’. Orthologous segments were identified based on flanking region synteny. We aligned all primate homologous SDs with MAFFT^57^ (v7.515) and used trimAl^58^ (v1.4) to remove the noise sequences with the parameter: ‘-automated1’. The phylogenetic trees were constructed with IQ-TREE^59^ (v2.1.4), and then we used BEAST^60^ (v2.6.6) with the HKY model incorporating gamma site, calibrated yule, and relaxed log-normal clock models to infer the split time. Node ages were estimated using log-normal priors. We conducted three independent runs for each tree and the results were consistent. Effective sample sizes exceeded 200 for all parameters in all runs.

For ILS analysis, we first truncated the fusion site flanking regions into 500 bp windows. Then, we utilized transanno (v0.4.5) (https://github.com/informationsea/transanno) to align these 500 bp segments to NHP genomes and generated multiple alignments across primate species. We then utilized IQ-TREE^59^ (v1.6.12) with HKY model and ete3 python package to analyze phylogeny trees, as described previously^44^.

### Subtelomeric repetitive region characterization, methylation analysis, and TE annotation

The pairwise identity heatmaps of each subtelomeric region were generated by ModDotPlot^61^ (https://github.com/marbl/ModDotPlot). We used SEDEF^62^ (v1.1r35) to annotate SDs. To analyze 5mC methylation levels in pCht regions and SD spacers, we utilized BEDTools^63^ (v2.30.0) to calculate mean methylation levels within 1 kbp windows, using a 500 bp step size. Synteny between subtelomeric repetitive regions was defined using minimap2 (v2.24) with the following parameters ‘-cx asm20 --secondary=no -A1 -B2 -O2,12 -s 1000 -Y -K 8G --eqx’. The syntenic relationship was visualized using R package circlize^64^ (v0.4.16). R package dendextend (v1.18.1)^65^ and TreeDist (v2.9.1)^66^ were utilized to plot the co-phylogeny and to calculate the generalized Robinson–Foulds distance.

### Centromere analysis

To analyze the structure of each chromosome, we first ran RepeatMasker (v4.1.4) on all chromosomes and identified the α-satellite-enriched regions as centromeres^67^.

Then, we utilized HumAS-HMMER (https://github.com/fedorrik/HumAS-

<u>HMMER_for_AnVIL</u>) to classify the suprachromosomal families (SFs) of α-satellites in human, *Pan*, gorilla, and orangutan. For macaque, the OWM-SF annotation tool was used to annotate the SFs^37^. The HOR arrays were characterized using the tool StV (https://github.com/fedorrik/stv).

However, for orangutan chr2a and chr2b, the tool encountered difficulties in correctly identifying the HORs, likely due to the complex structure of these acrocentric chromosomes. Thus, we selected the largest continuous regions containing the same SFs as their HORs. We then estimated the frequency of 5mC and CpG methylation within the HOR regions. BEDTools (v2.30.0) was used to count the frequency of methylation within 5 kbp windows and we determined the regions with the minimum frequency and below the lower quartile among the whole HOR as CDRs. Additionally, we used the previous published tool^68^ (https://github.com/altemose/chm13_hsat) to identity the HSatII/HSatIII arrays in human and *Pan* peri/centromeric regions (α-satellite-enriched regions and 5 Mbp on the p-arm and q-arm).

We further characterized the human centromere degenerate site through synteny comparisons between *Pan* and human, incorporating RepeatMasker annotations of ‘ALR_Alpha.’ Breakpoints between chimpanzees and humans were identified through alignment and analysis of specific SF organizations. ModDotPlot (https://github.com/marbl/ModDotPlot) was used to generate heatmaps for chimpanzee chr2b with a window size of 5,000 bp and for the human centromere degenerate site with a window size of 200 bp.

To assess the diversity of centromere degeneration across human populations, we first confirmed the presence of this site in all human genomes using minimap2 (v2.24) with the parameters ‘-cx asm20 --secondary=no -s 2500’. We then extracted the targeted regions from each assembly from HPRC data and ran RepeatMasker (v4.1.4) on these regions. We defined three satellite arrays—cenD_1, cenD_2, and cenD_3—with subtypes determined by length variations.

To confirm the fusion occurred in archaic humans, we randomly selected two individuals from the five structural haplotypes (total 10 individuals) and five NHP genomes to run mrsFAST^69^ (v3.4.2) for identifying singly unique nucleotide *k*-mers (SUNKs) in modern humans.

Subsequently, we checked the counts of the SUNKs in archaic human genomes.

To compare the methylation frequencies among human chr2, α-satellite arrays at the degenerate centromeric site, and NHP centromeres, we chunk the HORs (excluding CDRs; for macaques, we used the whole α-satellite region, excluding CDRs) into 17.1 kbp windows by BEDTools (v2.30.0) with parameters: ‘-w 171000 -s 8550’ and calculate the frequencies within windows.

Visualizations were generated using ggplot2.

### Fusion site KO experiments and RNA-seq analysis

The design of optimal sgRNA pairs to target sites was performed using the online CRISPR design tool (http://crispor.gi.ucsc.edu/). The complementary oligonucleotide pairs of L-sgRNA #1/#2 and R-sgRNA were annealed at 95℃ for 5 min, with ramp-down to 25℃ to generate the double-stranded DNA (dsDNA) fragment, before ligation into BbsI-digested PX459 (plasmid #48139, Addgene).

HEK-293T cell lines were maintained in complete culture medium: DMEM (Dulbecco’s modified Eagle’s medium) supplemented with 10% FBS (fetal bovine serum), 1% Penicillin-Streptomycin Solution (100×stock, Gibco). Transfection of CRISPR plasmid DNA into HEK- 293T cells was performed using Lipofectamine^TM^ 3000 Reagent (Invitrogen). Briefly, 0.5– 5×10^5^/cm^2^ HEK293T cells in 6 well plates were transfected with 2.5 μg of CRISPR plasmid DNA. Then, 24 hours after transfection, selection was conducted by adding puromycin (1 μg/ml) to the media for the next 48 hr. Surviving cells were cultured for 4-7 days without selection before harvesting.

Genomic DNA (gDNA) was extracted from transfected HEK-293T from each subline (1×10^5^) with the TIANamp genomic DNA Kit (DP304-02; Tiangen) according to the manufacturer’s instructions. Genome deletions caused by the sgRNA pairs were detected by PCR-amplification of gDNA using a primer pair flanking the deletion. The primers used for PCR screening are listed in Supplementary Table 8. Genomic PCR products were detected by agarose gel electrophoresis and Sanger sequencing (Shanghai, TsingKe).

Total RNA of the fusion site depletion and wild-type cell lines were reverse-transcribed using HiScript II Q RT SuperMix for qPCR kit (Vazyme), respectively. Gene expression was normalized to *GADPH*, and fold changes were calculated as described in the Figure 5. Error bars in the RNA analysis represent the standard deviation of the average fold changes based on at least two cell lines and/or two experimental replicates as indicated in the Figure 5. The primer sequences are listed in Supplementary Table 12.

We utilized fastp^70^ (v0.23.2) to perform adaptor trimming of raw sequencing reads. The trimmed reads were aligned to the reference genome (T2T-CHM13v2.0) using hisat2^71^ (v2.2.1).

Differential expression analysis was conducted using the R package DESeq2^72^ and edgeR^73^ and we identified genes as significantly differentially expressed with following criteria: false discovery rate (FDR) < 0.05 and a log2 fold change > 2. The genes considered significantly differentially expressed met two criteria: they were identified by both tools and demonstrated significance in two independent sgRNA design experiments, forming the final set. Visualizations were generated using ggplot2. We performed Gene Ontology (GO) enrichment analysis using DAVID GO^74^ (https://david.ncifcrf.gov/).

## Acknowledgments

We thank Tonia Brown for editing this manuscript. We thank the HPRC and Primate T2T Consortium for providing the long-read human and great ape genome assemblies. The computations in this study were run on the Siyuan-1 supported by the Center for High Performance Computing at Shanghai Jiao Tong University. E.E.E. is an investigator of the Howard Hughes Medical Institute.

This article is subject to HHMI’s Open Access to Publications policy. HHMI lab heads have previously granted a nonexclusive CC BY 4.0 license to the public and a sublicensable license to HHMI in their research articles. Pursuant to those licenses, the author-accepted manuscript of this article can be made freely available under a CC BY 4.0 license immediately upon publication.

## Author contributions

Y.M., E.E.E., and Q.S. conceived the project; X.J., Z.Y., X.Y., K.M., S.Z., J.C., J.H., L.F., J.Z., and Y.M. contributed to the syntenic comparison, fusion site characterization, ILS, and subtelomeric repetitive region analyses; L.Z., X.J., Y.L., Y.N., X.B., and Q.S. contributed to the fusion site depletion analysis; D.Y. and E.E.E. generated the genome assemblies of great apes; X.J., G.Z., and Y.M. analyzed the centromeres. Y.M. and E.E.E. wrote the draft manuscript with contributions from other authors. All authors read and approved the manuscript.

## Data Availability

The T2T primate genomes used in this study are available from GenBank via accessions: GCA_009914755.4, GCA_028858775.2, GCA_028885625.2, GCA_028885655.2, GCA_029281585.2, GCA_029289425.2 and GCA_037993035.1. The T2T primate genome assemblies are also available on GitHub (https://github.com/marbl/Primates and https://github.com/zhang-shilong/T2T-MFA8). The Neanderthal and Denisovan genomes used are available from https://www.eva.mpg.de/genetics/genome-projects. The Iso-Seq data are available on ENCODE (tissues: ENCFF492BYP, ENCFF306ZPP, ENCFF318SKH, CHM13- T2T2 hTERT Iso-Seq: SRR12519035, SRR12519036).

